# Scalable distance-based phylogeny inference using divide-and-conquer

**DOI:** 10.1101/2023.10.11.561902

**Authors:** Lars Arvestad

## Abstract

Distance-based methods for inferring evolutionary trees are important subroutines in computational biology, sometimes as a first step in a statistically more robust phylogenetic method. The most popular method is Neighbor Joining, mainly to to its relatively good accuracy, but Neighbor Joining has a cubic time complexity, which limits its applicability on larger datasets. Similar but faster algorithms have been suggested, but the overall time complexity remains essentially cubic as long as the input is a distance matrix. This paper investigates a randomized divide-and-conquer heuristic, `dnctree`, which selectively estimates pairwise sequence distances and infers a tree by connecting increasingly large subtrees. The divide-and-conquer approach avoids computing all pairwise distances and thereby saves both time and memory. The time complexity is at worst quadratic, and seems to scale like *O*(*n* lg *n*) on average. A simple Python implementation, `dnctree`, available on GitHub and PyPI.org, has been tested and we show that it is a scalable solution. In fact, it is applicable to very large datasets even as plain Python program.

## 1 Introduction

Although distance-based methods for phylogeny inference have been dismissed for a long time due to low accuracy, they are still relevant in Computational Biology. Statistical methods are preferred when a correct phylogenetic tree is the main goal, but distance-based methods are important as subroutines to other methods thanks to ease of use, ease of implementation, and good-enough speed in many applications. For example, statistical phylogeny inference may need a starting tree and tools for multiple sequence alignment often use a rapidly computed guide tree. Even quite imperfect trees may yield good end-results for such applications.

Neighbor Joining (NJ), by Saitou and Nei (1987), has become the most popular distance-based method and its benefits have been studied both empirically and theoretically. For example, NJ has a convergence radius, which guarantees that the correct tree is returned if the largest error on a pairwise distance is *δ/*2 when the shortest branchlength is *δ* in the tree that generated the sequences (Atteson, 1999). A more general result is given by Mihaescu et al. (2009), where it is also shown that NJ remains correct far beyond Atteson’s guarantee. The appealing quality output of NJ is however often overshadowed by its cubic time complexity, rendering it challenging to use on even medium-sized datasets by modern standards.

A number of strategies have been suggested for NJ or NJ-like inference on large datasets. Good programming and careful optimization of “classic” NJ can push the practical limits further. Wang et al. (2023) implemented NJ using vectorization, threading using OpenMP, and branch-and-bound techniques from RapidNJ to make it possible to handle 64,000 sequences.

For true scalability, algorithmic improvement is needed. Elias and Lagergren (2009) suggested FNJ, an NJ-like algorithm using the same selection function, that avoids evaluating all vertex pairs in every iteration. Instead, information is retained from iteration to iteration for choosing a partner for each newly created node, so that the next NJ-like pairing is computed in *O*(*n*) time. While this sacrifices some correctness on noisy datasets, the time complexity is quadratic. The algorithm has Atteson’s convergence radius and therefore has the same quality guarantee as NJ. Khan et al. (2013) showed that using a careful implementation of pairwise distance estimation and applying FNJ can manage 10^5^ sequences at a reasonable time, but needing about 25 GiB RAM, mainly for storing the input distance matrix.

FastTree (Price et al., 2009) uses ideas from FNJ and combines with other heuristics, including comparing alignment profiles rather than sequences. Using a greedy postprocessing improvement step, nice accuracy is achieved while retaining fast computation. A maximum-likelihood heuristic was later used in the postprocessing (Price et al., 2010).

Simonsen et al. (2008) proposed the NJ variant RapidNJ which utilizes different search strategies and a bounding technique to avoid bad choices of node pairings. While the worst-case time is *O*(*n*^3^), the authors show that there is a significant speedup on most data. Indeed, it has been shown to manage inputs with 10^5^ sequences (Simonsen et al., 2008; Khan et al., 2013).

Clausen (2023) suggested two heuristics, DNJ and HNJ, based on operations similar to FNJ, and carefully engineered implementations that are shown to handle a distance matrix for 10^6^ input elements in about a day on a modern computer with 512 GB of RAM. Clausen notes that memory, not time, becomes the bottleneck with quadratic time algorithms.

Providing fast NJ-like algorithms is however not the only step needed for scalable phylogeny inference, as Elias and Lagergren (2007) pointed out. Even if tree inference on *n* sequences is *O*(*n*^2^), the input is a distance matrix that needs 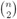 distances estimated and if their average length is *L*, then the time complexity of tree inference needs *O*(*n*^2^*L*) time — still essentially cubic time complexity.

Kannan et al. (1996) realized early the importance of avoiding to compute pairwise distances and suggested an algorithm that used *O*(*n* lg *n*) experiments to iteratively place leaves into the expanding tree. One experiment decides, for three sequences, which rooted tree best describes their evolution. Using a balanced search tree, they ensure that a leaf insertion is essentially performing a binary search to determine which edge to attach to. Lingas et al. (2001) suggested a divide-and-conquer method, also based on using rooted triples to determine where locations, where the leaf set is partitioned into two subsets on which two trees are computed and subsequently merged. The authors note that the method extends to unrooted trees using quartets, in which one leaf is chosen as an outgroup. Brodal et al. (2001) continued this work and designed a datastructure to support building *d*-degree rooted trees in *O*(*n* lg_*d*_ *n*).

We suggest a divide-and-conquer method that takes a set *S* of sequences of length *L* and a distance-estimation method 𝒟: *S* × *S* → ℝ^+^ as input, and compute an unrooted tree with elements of *S* labeling the leaves. The method carefully chooses quartets that partition *S* into three subproblems, and these quartets determines which distances to compute. The resulting trees from the subproblems are connected to form a resulting tree. Importantly, the suggested algorithm has Atteson’s convergence radius, thus guaranteeing correct phylogenies when yields good results. The base case is computed using NJ, and the larger base case one can afford, the better the quality of the output.

The algorithm has been implemented in a simple Python program, `dnctree`, which is easily installed from PyPI.org. Source code is available at https://github.com/arvestad/dnctree. Basic performance tests and a comparison with NJ are presented.

## 2 Methods

### 2.1 The dnctree algorithm

The algorithm idea is to let three randomly chosen sequences guide a partitioning of the input into three subproblems. With a carefully chosen partitioning, constructing a joint tree becomes simple. The base case are datasets with less than *K* sequences, in which case NJ is used.

The `dnctree` algorithm takes a set of sequences *V* and a distance function *𝒟* as input and can be summarized as follows.

1. If |*V* | ≤ *K*:
  A. Let *M* be all pairwise distances for *V*, computed using. 𝒟
  B. Return NJ(*M*).
2. Pick three sequences *x, y, z* ∈ *V* at random.
3. Estimate distances *d*_*xy*_, *d*_*xz*_, and *d*_*yz*_ using 𝒟.
4. Let *c* be a “center” vertex and choose *d*_*xc*_, *d*_*yc*_, *d*_*zc*_ optimally.
5. Create subsets *V*_*x*_ = {*x, c*}, *V*_*y*_ = {*y, c*}, and *V*_*z*_ = {*z, c*}
6. For υ ∈ *V* \ {*x, y, z*}:
  A. For *w* ∈ {*x, y, z*}: *d* _υ *w*_ = 𝒟 (υ, *w*)
  B. Compute a quartet test on {*x, y, z*, υ}.
  C. Place υ accordingly: in *V*_*x*_, *V*_*y*_, or *V*_*z*_.
  D. If υ ∈ *V*_*x*_, set *d*_υ*c*_ = (*d*_υ*y*_ + *d*_υ*z*_ *-d*_*yz*_)*/*2, or correspondingly when υ∈ *V*_*y*_ or υ ∈ *V*_*z*_.
7. For *i* ∈ {*x, y, z*}: *T*_*i*_ = `dnctree`(*V*_*i*_, *D*)
8. Return *T* = *T*_*x*_ ∪ *T*_*y*_ ∪ *T*_*z*_

Step 4 defines a tree *T*_*xyz*_ on *x, y* and *z* with *c* in the center. The branch-lengths of *T*_*xyz*_ satisfies

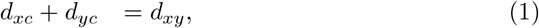

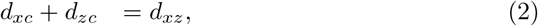

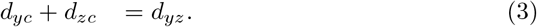

Step 6c uses a quartet test to determine which of *x, y* and *z* that the other sequences are most closely related to, in way determining in which part of *T*_*xyz*_ they ought to be placed.

In step 8 a new tree is created by connecting *T*_*x*_, *T*_*y*_, and *T*_*z*_ on *c*. Distances are recomputed, according to the presentation above, but can and should be cached. Allocating a distance matrix is wasteful, except in the base cases, because most entries are zero when *K* ≪ | *V*. |

Ideally, we get three subproblems of roughly equal size, which would yield time complexity *O*(*n* lg *n*). Input data may not support a tree with that property, but we are guaranteed that the largest subproblem has size *n* - 2, which implies that there are at most *O*(*n*) iterations, and thus *O*(*n*^2^) steps.

### 2.2 Implementation and hardware

Both `dnctree` and NJ was implemented anew in Python (single threaded). While NJ might be unnecessary to implement again^1^, it was needed as an easy-to-use subroutine for `dnctree`.

All testing was done on a MacBook Pro with an Apple M1 Pro CPU and 16 GB RAM. Run time was measured with the `time` command using the “user” field.

### 2.3 Data

Simulated test data was generated by constructing symmetric additive trees with 127 leaves (see Figure 1a). Two innermost branches were set to length 1.0, the rest has length 0.1. For each parameter combination there was 100 replicates.

**Figure 1.**
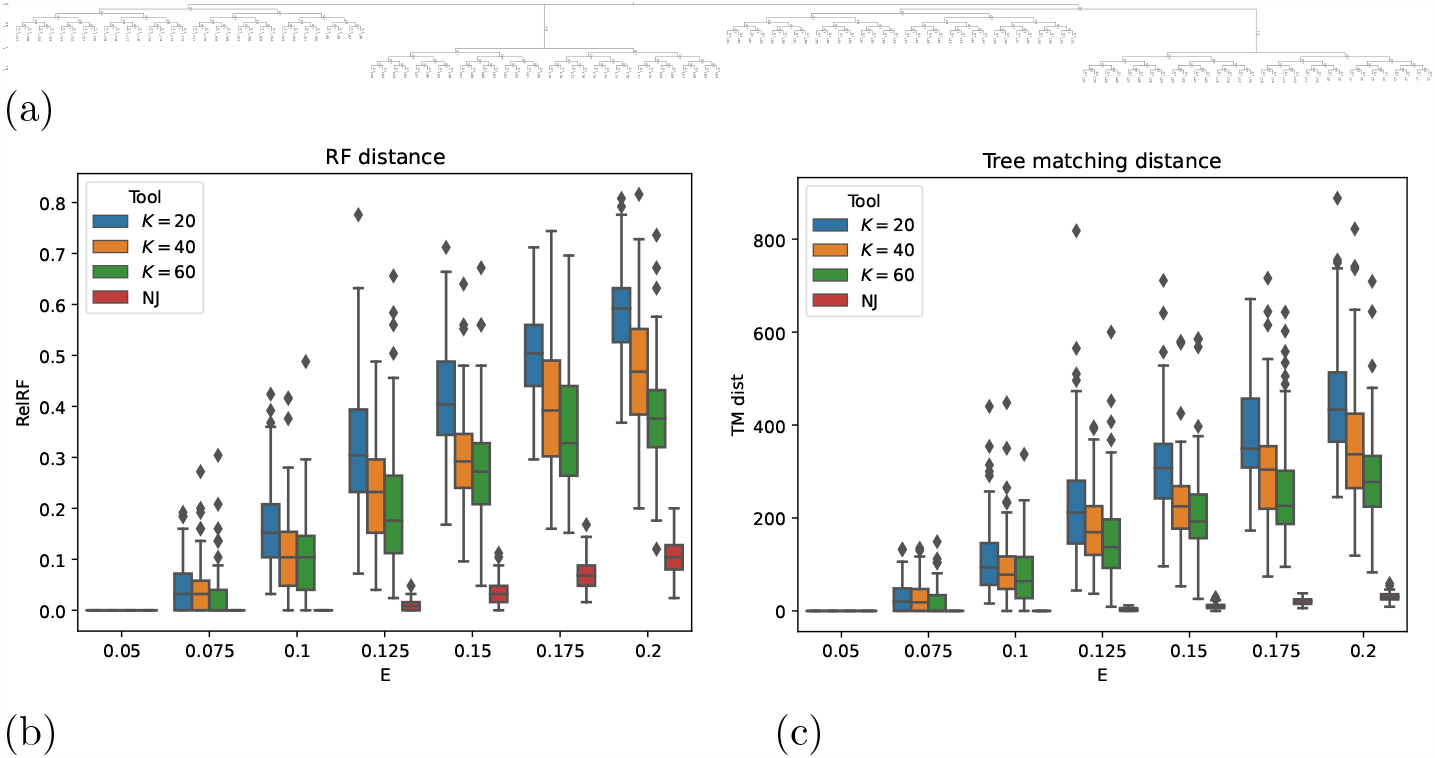
Tests on simulated distances (no sequences generated). *K* is the base case size when running `dnctree` and *E* determines the error distribution: uniformly on [- *E, E*]. The results are plotted using seaborn (Waskom, 2021) box plots, indicating quartiles (minus outliers as determined by seaborn). For *E* = 0.05, Atteson’s convergence radius is corroborated: all trees are recovered perfectly. (a) The choice of 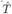 to which noise was added. Two edges has length 1.0 and the others 0.1. (b) Results assessed with relative Robinson-Foulds distances. (c) Results assessed with TMD.

For testing without sequences, noise was added to the additive distances by sampling error terms from [- *E, E*], with *E* being a parameter varied in testing (see Figure 1).

When testing with sequence data, AliSim (Ly-Trong et al., 2022) was run using the WAG model (Whelan and Goldman, 2001) and otherwise default parameters. Alignments had widths 500 and 1000 residues, in different experiments.

For testing on biological data, some Pfam (Mistry et al., 2021) alignments were downloaded from InterPro (Paysan-Lafosse et al., 2023). The main criteria was that the Pfam alignments contained many sequences and many alignment columns. The chosen Pfam alignments were subsampled to 2^*k*^ × 1000 sequences, for *k* from 0 to 7.

### 2.4 Evaluating inference accuracy

We assessed inferred tree accuracy with two functions. The relative Robinson-Foulds distance (Robinson and Foulds, 1981) has the advantage that it used by many authors and the noted disadvantage of being sensitive to small errors (Bryant and Steel, 2009). It was computed using ETE3 (Huerta-Cepas et al., 2016).

The Tree Matching Distance (TMD, by Lin et al., 2011) was used as a complement due to being reported as harder to saturate, and computed using the Python package `tree-matching-distance`, written by this author and available on PyPI.org.

## 3 Results

### 3.1 Atteson’s convergence radius

Let 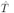 be the tree that originally displayed the sequences of *S*, and let δ be the shortest branch of *T*. Let *D* be the pairwise distance matrix on *S* computed using 𝒟. Atteson (1999) showed that, if max_*a,b*∈*S*_ 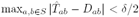 < δ/2, then NJ with *D* as input returns 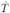. Due to how `dnctree` uses NJ as a subroutine, it also benefits from Atteson’s guarantee.

#### Theorem 1.

*If the largest error in D is δ/*2, *then* `dnctree` *will return* 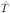

*Proof*. We use induction and follow the outline of the algorithm in 2.1. If *S* is at most *K*, then NJ is utilized and Atteson’s guarantee holds. We turn to larger inputs and assume that `dnctree` is correct for inputs of size *m* or smaller, and consider size *n* = *m* + 1. There are two algorithm parts: partitioning in step 6 and the recursive calls. Let 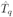 be the restriction of 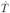 to a quartet *q*_*v*_ ={ *x, y, z, v* }computed to determine a subset for *v*. The smallest edge of 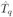 is at most δ and the largest error from 𝒟 is still δ*/*2. Therefore, the Atteson bound guarantees that 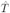 is inferred on *q*_*v*_ and *v* will be correctly assigned to a subset. This holds for all *v*, so the partitioning step is correct.

In the worst case, the largest subset is *S* { *c* } \ { *x, y, z* } and therefore of size at most *n* − 1, which by the induction hypothesis means that the three recursive calls in step 7 will return the correct subtree. A merged subtree (line 8) on the center vertex *c*, which corresponds to the vertex joining *x, y*, and *z* in all quartets, which means that the connected result is 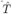.

### 3.2 Testing with simulated distances

In a first test, a distance matrix was initialized using distances from a given tree with a noise term added. The noise was sampled from a uniform distribution on [- *E, E*].

Both NJ and `dnctree`, for all choices of *K* recovers all trees perfectly when *E* = 0.05, which is to be expected with a correct implementation because the smallest branch in the true tree is 0.1 and Atteson’s convergence radius guarantees correct inference.

Inference suffers when noise increases and `dnctree` performs poorly compared to NJ. This is expected since `dnctree` uses less information when making decisions on branching. As expected, base case size *K* affects inference quality and there is a clear beneficial effect of having large base cases. NJ manages remarkably well regardless of errors being outside the convergence radius, which has been noted before (Atteson, 1999; Mihaescu et al., 2009; Gascuel and Steel, 2016).

Other choices of simulated trees yielded similar results.

### 3.3 Testing with simulated sequence alignments

For simulated alignments, trees were simulated in a Yule-Harding process and then sequences were evolved in those trees (Ly-Trong et al., 2022). As expected, the quality of the inferred trees varies a lot, see Figure 2, but larger base cases and more data is beneficial. NJ has clearly more accurate inference, but decent distance estimates can still generate viable inference quality with `dnctree`.

**Figure 2.**
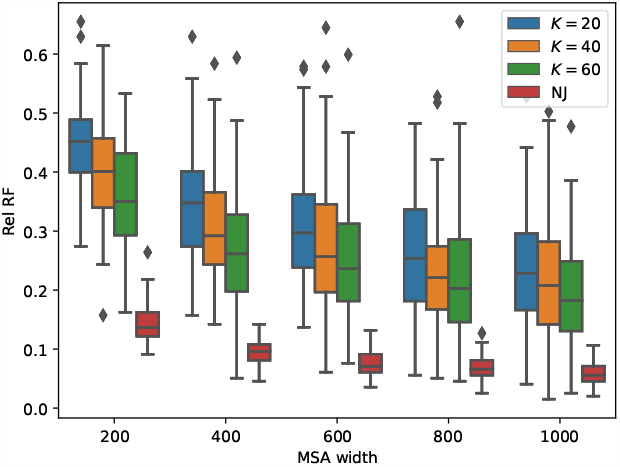
Relative Robinson-Foulds distance on simulated alignments of different alignment widths and different base case sizes *K*. There are 100 replicates for each parameter combination. Each alignment was generated in a tree with 200 leaves and the aligment width is 1000 residues.

The same pattern is seen in a larger experiment, pushing to quite large datasets of up to 3000 leaves and a MSA width of 500 columns. Figure 3a shows that `dnctree` scales almost linearly, there is just a hint of the log-factor at tested datasizes, when looking at the number of computed pairwise distances, which is the bottleneck in the experiment.

**Figure 3.**
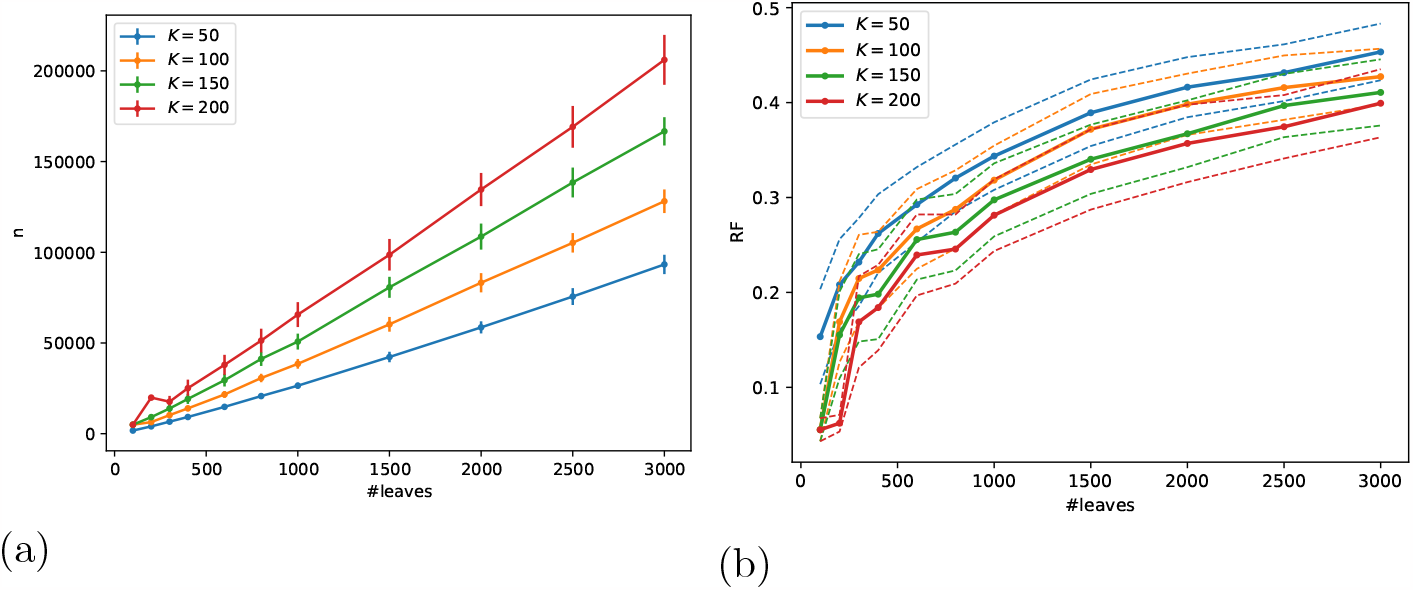
Work and accuracy. Random trees and sequences were generated with AliSim. (a) Number of pairwise distances computed at different dataset sizes. Vertical bars indicate standard error. (b) The mean accuracy for each base case size, taken over 100 replicates per parameter combination, is indicated with a solid line and the dotted lines with same color indicate the standard error.

### 3.4 Testing on biological data

To demonstrate the utility of `dnctree`, we ran some tests with subsampled Pfam alignments, see Figure 4. The tests indicate that `dnctree` scales as expected from the tests on simulated data. Runtime is increasing almost linearly with input size, regardless of dataset, and there is some variation due to the underlying phylogeny and choice of sequence triples that partition the subproblems.

**Figure 4.**
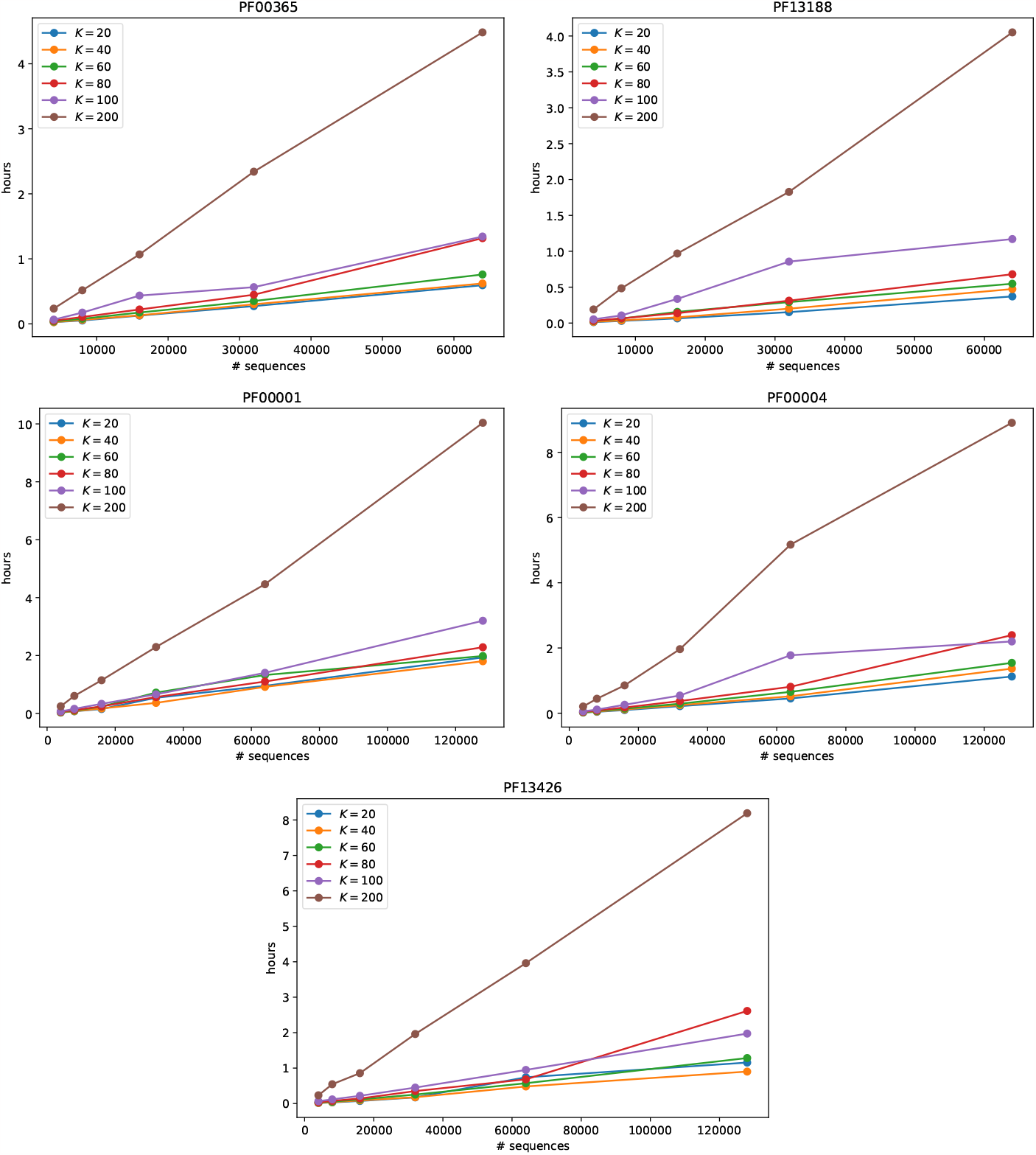
Scalability tests on Pfam data. The lines are for different base case sizes *K*. Sequences from the Pfam alignments were subsampled with up to 64,000 (PF00365, PF13188) or 128,000 sequences (PF0001, PF0004, PF13426). The runtime in hours on the vertical axis. The choice of *K* does not fully determine runtime, as is shown by lines mixing somewhat. This is due to the random choices of sequence triples that determine subproblems, that are not always ideal.

The observed runtimes are practical even on the largest datasets. A phylogeny can be inferred on challenging datasizes in a few hours, even with a pure-Python implementation running on a regular laptop.

## 4 Discussion

The tested implementation of `dnctree` is written in pure Python, without any kind of clever speedups, and yet it was possible to infer evolutionary trees for large datasets on rather simple hardware. Since a tiny fraction of all pairwise distances are actually computed, RAM is not a big concern. The quality of the inferred trees are surely very low, but they still have utility as, for example, starting points for tree exploration by a maximum-likelihood inference or as a rough first clustering of sequences.

As noted by Clausen (2023), memory is currently the bottleneck when using distance-based methods. However, since `dnctree` exhibits a *n* lg *n* memory usage in our testing (see Figure 3a), large datasets become feasible targets also on simple hardware. A dataset with 128,000 sequences and a base case size of *K* = 200 needs about 1 GB of RAM in our non-optimized Python implementation. Extrapolating based on our experiments suggests that datasets with 10^6^ sequences can be managed on common hardware in a couple of days. Encouragingly, preliminary experiments suggests that the possible speedup is two orders of magnitude if implementing `dnctree` in for example C++, and memory usage is likely to be lower as well.

The focus of our testing has been on showing that the method can be applied to very large datasets with very small computational resources. Therefore, the base case size has been quite small, trading output accuracy for speed. For serious use cases, one would push the base case size as high as one can afford. The base cases are an advantage with the propsed heuristic: you can and should choose large base cases. The way the data is partitioned, base cases tend to be for “natural” subtrees, which is also why the merging step becomes very easy to implement.

A phylogeny heuristic that drastically reduces the number of pairwise distances computed opens up for more computational heavy distance estimation. Bogusz and Whelan (2017) experimented with a method that integrated over all possible alignments of two sequences, using a pair-HMM that models pair-wise alignments. They showed that for simulated sequences the estimated distances were closer to additive than distances estimated from multiple sequences alignments (MSAs). Combining their distance estimator with NJ was rivaling RAxML (Stamatakis, 2014) applied to MSAs computed with MAFFT (Katoh and Standley, 2013). Using the method by Bogusz and Whelan in `dnctree` might improve how subproblems are partitioned, a sensitive step in `dnctree`, and thus improve overall accuracy. At the same time, `dnctree` would extend the applicability of their distance inference method to much larger input instances than was conceivable with NJ.

## 5 Availability

Source code for `dnctree` is available at https://github.com/arvestad/dnctree. Installing from PyPI.org (run the shell command pip install dnctree; see https://pypi.org/project/dnctree/) is recommended for most users.

## Footnotes

1 It must be one the of the most commonly implemented algorithms in the world.

